# Post-mitotic expansion of cell nuclei requires ACTN4-mediated nuclear actin filament bundling

**DOI:** 10.1101/2020.04.29.067488

**Authors:** Sylvia Krippner, Jannik Winkelmeier, Carsten Schwan, Julian Knerr, David Virant, Ulrike Endesfelder, Robert Grosse

## Abstract

The actin cytoskeleton operates in a multitude of cellular processes including cell shape and migration, mechanoregulation, as well as membrane or organelle dynamics. However, its filamentous properties and functions inside the mammalian cell nucleus are less well explored. We previously described transient actin assembly at mitotic exit that promotes nuclear expansion during chromatin decondensation. Here, we identify non-muscle ACTN4 as a critical regulator to facilitate F-actin formation, reorganization and bundling during postmitotic nuclear expansion. ACTN4 binds to nuclear actin filaments and ACTN4 clusters associate with nuclear F-actin in a highly dynamic fashion. ACTN4 but not ACTN1 is required for proper postmitotic nuclear volume expansion, mediated by its actin binding domain. Using super-resolution imaging to quantify actin filament numbers and widths in individual nuclei we find that ACTN4 is necessary for postmitotic nuclear actin assembly and actin filament bundling. Our findings uncover a nuclear cytoskeletal function for ACTN4 to control nuclear size during mitotic cell division.

## Introduction

The actin cytoskeleton fulfils many essential cellular functions such as membrane protrusion and adhesion, contractility and cytokinesis, polarity and molecular transport. These functions are executed largely through its ability to dynamically assemble and disassemble filamentous actin (F-actin). To achieve this, the actin cytoskeleton is controlled through a plethora of actin binding and regulatory proteins including factors that promote structural organization for instance through F-actin crosslinking or bundling (Chhabra and Higgs, 2007; Lee and Dominguez, 2010).

Although much is known about the dynamics and functions of cellular F-actin structures, their roles and properties in the nuclear compartment of living cells remain largely unexplored (Plessner and Grosse, 2019). Photobleaching experiments of GFP-actin suggested the existence of some dynamic equilibrium (McDonald et al., 2006). Recent tools allowing for the direct visualization of nuclear actin assembly in real time have enabled significant insight into the characteristics and functions of the actin cytoskeleton inside the nucleus (Melak et al., 2017). Nuclear actin filaments have thus been directly observed and monitored in live mammalian cells and implicated for example in rapid signaling to transcriptional or chromatin dynamics (Baarlink et al., 2013; Tsopoulidis et al., 2019; Wang et al., 2019), in homology-directed DNA repair (Schrank et al., 2018) and myosin-dependent relocalization of heterochromatin breaks (Caridi et al., 2019), in DNA replication (Parisis et al., 2017) or in virus-induced nuclear envelope disruption and egress of viral particles (Ohkawa and Welch, 2018).

We have previously shown that a dynamic nuclear actin cytoskeleton forms and reorganizes during expansion of daughter cell nuclei after the exit of mitosis (Baarlink et al., 2017). In mammalian cells, nuclear volume expansion after mitosis occurs throughout early G1 when chromatin becomes decondensed to reestablish a functional G1 interphase nucleus, a highly complex process involving nuclear envelope reassembly and reorganization of nuclear architecture (de Castro et al., 2016; Gerlich et al., 2001; Schooley et al., 2012; Webster et al., 2009). We could show that the actin severing and disassembly factor Cofilin-1 plays a pivotal role in controlling the timing and turnover dynamics of nuclear actin filaments during mitotic exit and nuclear growth (Baarlink et al., 2017). However, how these filaments are organized and assembled or whether other actin-regulating proteins function during this process was unknown (Moore and Vartiainen, 2017).

## Results and Discussion

We previously established a phalloidin-based nuclear capture assay to identify proteins interacting with endogenous nuclear actin filaments at mitotic exit and identified Cofilin-1 and its nuclear function in cell division (Baarlink et al., 2017). In addition, this approach revealed non-muscle alpha-actinin 4 (ACTN4) as a top candidate binding nuclear F-actin at mitotic exit (Baarlink et al., 2017). ACTN4 and its close homologue ACTN1 are spectrin repeat (SR)-containing proteins forming antiparallel homodimers, thereby facilitating interactions with F-actin via their N-terminal actin-binding domain (ABD) (Fig. 1A) and have been implicated in cell adhesion, motility, proliferation and cancer progression (Honda, 2015). ACTN4 is known as a potent actin bundling and crosslinking factor (Courson and Rock, 2010; Honda et al., 1998; Hotulainen and Lappalainen, 2006; Travers et al., 2013; Winkelman et al., 2016). Notably, ACTN4 was shown to shuttle between the cytoplasm and the nucleus in a CRM1-dependent manner (Kumeta et al., 2010). We therefore set out to investigate its potential role for nuclear actin assembly and reorganization during mitotic exit.

**Figure 1.**
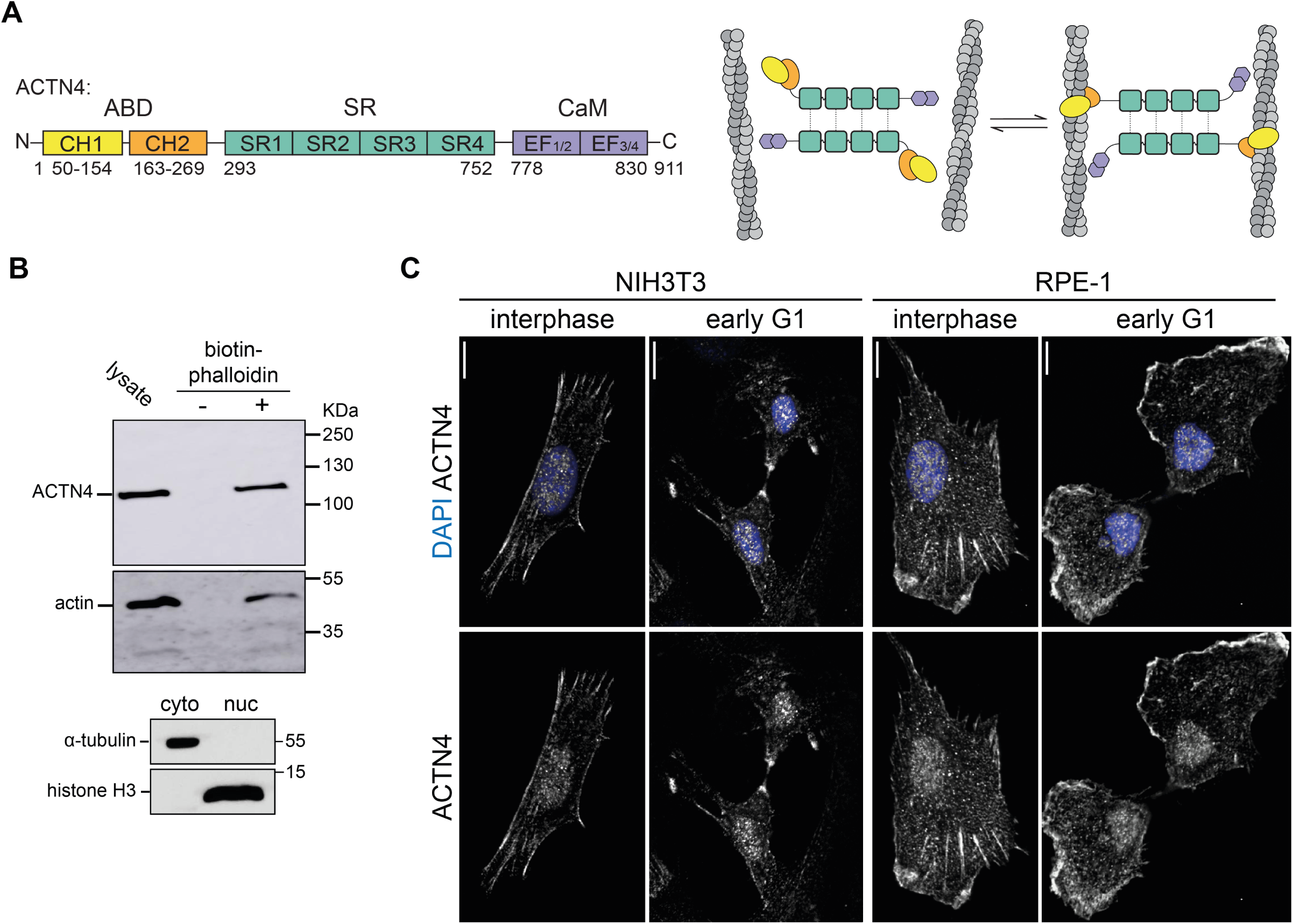
ACTN4 interacts with postmitotic nuclear actin filaments and localizes to the nucleus. **(A)** Domain architecture and crosslinking function of ACTN4. actin-binding-domain (ABD), calponin-homology (CH1, CH2), spectrin-repeats (SR), calmodulin-like domain (CaM), EF-Hand domains. **(B)** Upper panel: Immunoblot showing precipitation of ACTN4 and β-actin from RPE-1 nuclear lysates in early-G1 in the presence or absence of Biotin-Phalloidin as indicated. Lower panel: Immunoblot of cytosolic-(cyto) and nuclear (nuc) fractions as indicated **(C)** Immmunofluorescence images of NIH3T3 or RPE-1 cells in interphase or early-G1. Cells were fixed and stained as indicated. ACTN4 was labelled by AF488 secondary antibody and DNA was stained with DAPI; scale bars 10 µm.

In order to validate ACTN4 as a nuclear F-actin binding partner in the early G1 phase of the cell cycle, we performed biotin-phalloidin pulldown assays. For this, RPE-1 cells were synchronized with the Cdk1 inhibitor RO-3306 at the G2/M border before washout after 2h to enrich cells in early G1. Cells were then subjected to subcellular fractionation to obtain nuclear extracts as determined by the presence of histone H3 and the absence of α-tubulin. Immunoblotting revealed that endogenous ACTN4 was present in nuclear fractions and that it was able to interact with endogenous nuclear F-actin (Fig. 1B). Consistent with this, we observed endogenous localization of ACTN4 to the nucleus of NIH3T3 or RPE-1 cells by immunostaining at mitotic exit but also in interphase cells (Fig. 1C), confirming previous results showing nucleocytoplasmic shuttling of ACTN4 (Kumeta et al., 2010). No obvious difference could be observed in nuclear localization of ACTN4 at early G1 or in interphase cells (Fig. 1C), indicating continuous nuclear distribution or shuttling and no specific nuclear accumulation after mitosis.

Since we observed interaction of ACTN4 with F-actin in nuclear fractions, we wanted to assess whether ACTN4 would colocalize with nuclear actin structures during mitotic exit. For this we used a genetically encoded nuclear-targeted anti-actin nanobody (nAC) as previously described (Plessner et al., 2015). This allowed for the visualization of endogenous nuclear actin together with SNAP-tagged ACTN4 and to follow their distribution and dynamics over time after spontaneous cell division during early G1 of NIH3T3 cells. This revealed the appearance of filamentous actin structures that were partially decorated by ACTN4 (Fig. 2A and B). Interestingly, ACTN4 appeared to dynamically slide along actin filaments but also localized to actin structures that seemed to be crosslinking events (Fig. 2A, Movie 1). Mobile ACTN4 clusters were visible over an extended period of time and allowed for automated tracking using Imaris (Fig. 2C). This shows that ACTN4 dynamically interacts with actin filaments during F-actin assembly and reorganization during early G1 of the cell cycle, suggesting that ACTN4 may play an important role in nuclear reorganization after mitosis.

**Figure 2.**
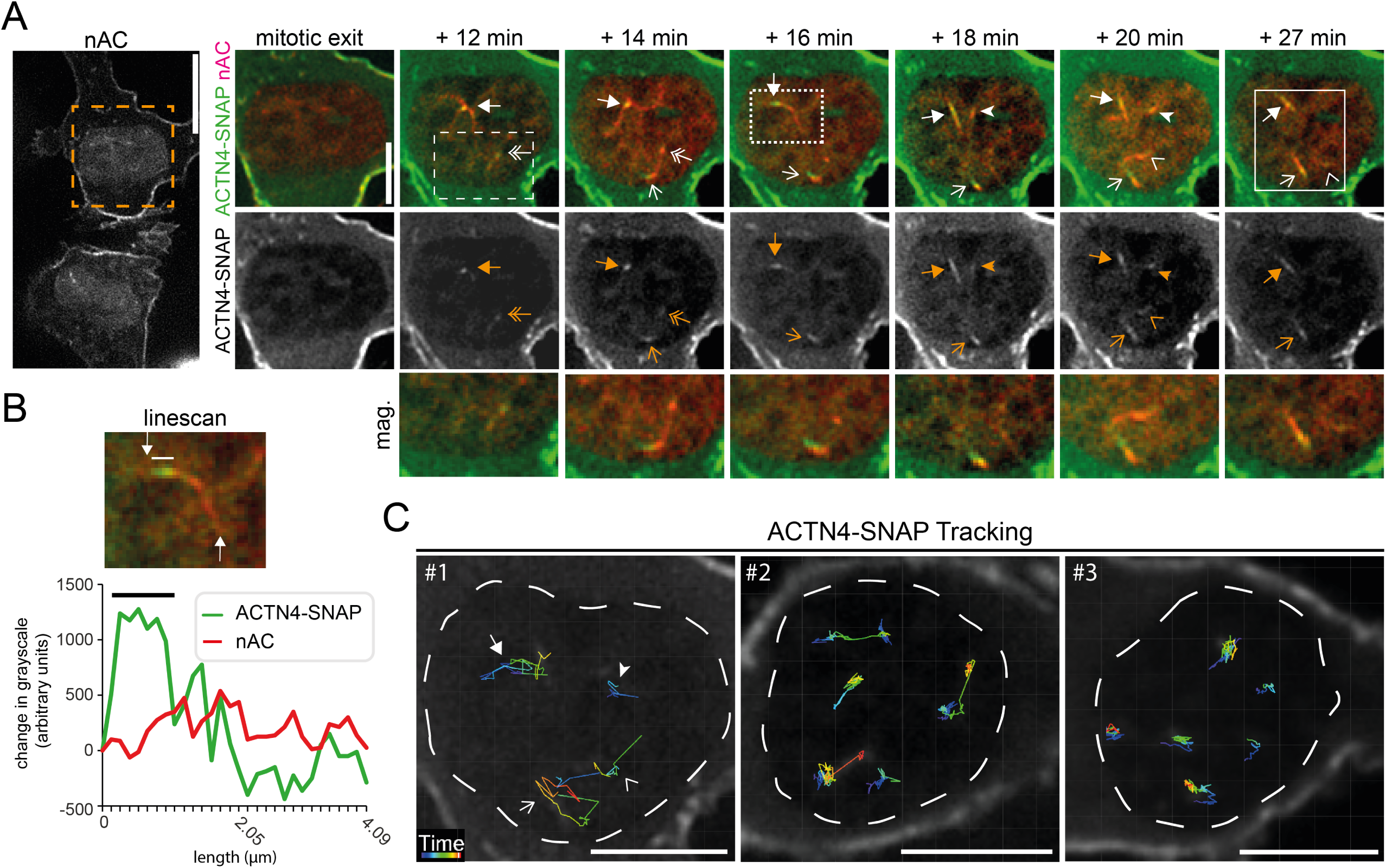
ACTN4 clusters dynamically associate with postmitotic nuclear actin filaments. **(A)** Cells in early G1 stably expressing nAC-mCherry and doxycycline-inducible ACTN4-snap were analyzed by time-lapse microscopy. ACTN4-snap was labeled by SiR 647 (green). Different arrows label dynamic ACTN4 clusters. The dashed white box is shown as magnification below. The solid box shows an actin filament with associated ACTN4 cluster that was analyzed by linescan (Fig 2B). Scale bar overview (nAC) represents 10 µm. Scale bar time series 5µm. **(B)** Linescan of an actin filament with associated ACTN4 cluster from the solid box in A. Arrows mark the ends of the filament subjected to linescan. A black-(linescan) or white (image) line highlights the ACTN4 cluster with increased intensity. **(C)** Automated tracking (autoregressive motion) of ACTN4 clusters in nuclei by IMARIS software. The nuclei are indicated by a dashed line. Tracks are visualized by colored lines. The color code reflects the time when an object was tracked (sequence #1: 27 min; sequence #2: 20 min; sequence #3: 15 min). Scale bars are 5 µm. Sequence #1 is also part of Fig. 2A. The different arrows represent clusters labeled in Fig. 2A.

We next wanted to examine whether ACTN4 is involved in nuclear actin-mediated expansion of daughter cell nuclei after mitosis. We therefore monitored actin assembly at mitotic exit by stably expressing nAC-mCherry in living NIH3T3 cells (Fig. 3A, Movie 2). As previously shown for nAC-tagGFP (Baarlink et al., 2017), we could observe the transient and dynamic formation and reorganization of F-actin structures in expanding daughter cell nuclei that spontaneously disassemble after approximately 90 minutes (Fig. 3A and B). As an initial loss of function approach, we performed double knockdowns of both, non-muscle isoforms ACTN1 and 4, using RNAi in NIH3T3 and RPE-1 cells expressing H2B-mCherry to visualize nuclei. Z-stacks of nuclei were obtained every 5 minutes for 90 minutes after mitotic exit. 3D reconstructed nuclear surfaces and corresponding volume measurements were obtained using Imaris. Notably, double knockdown of ACTN1 and ACTN4 in NIH3T3 resulted in a significant decrease in nuclear volume within 90 minutes after mitotic exit (Fig 3B), which was slightly more pronounced in RPE-1 cells (Fig. 3C).

**Figure 3.**
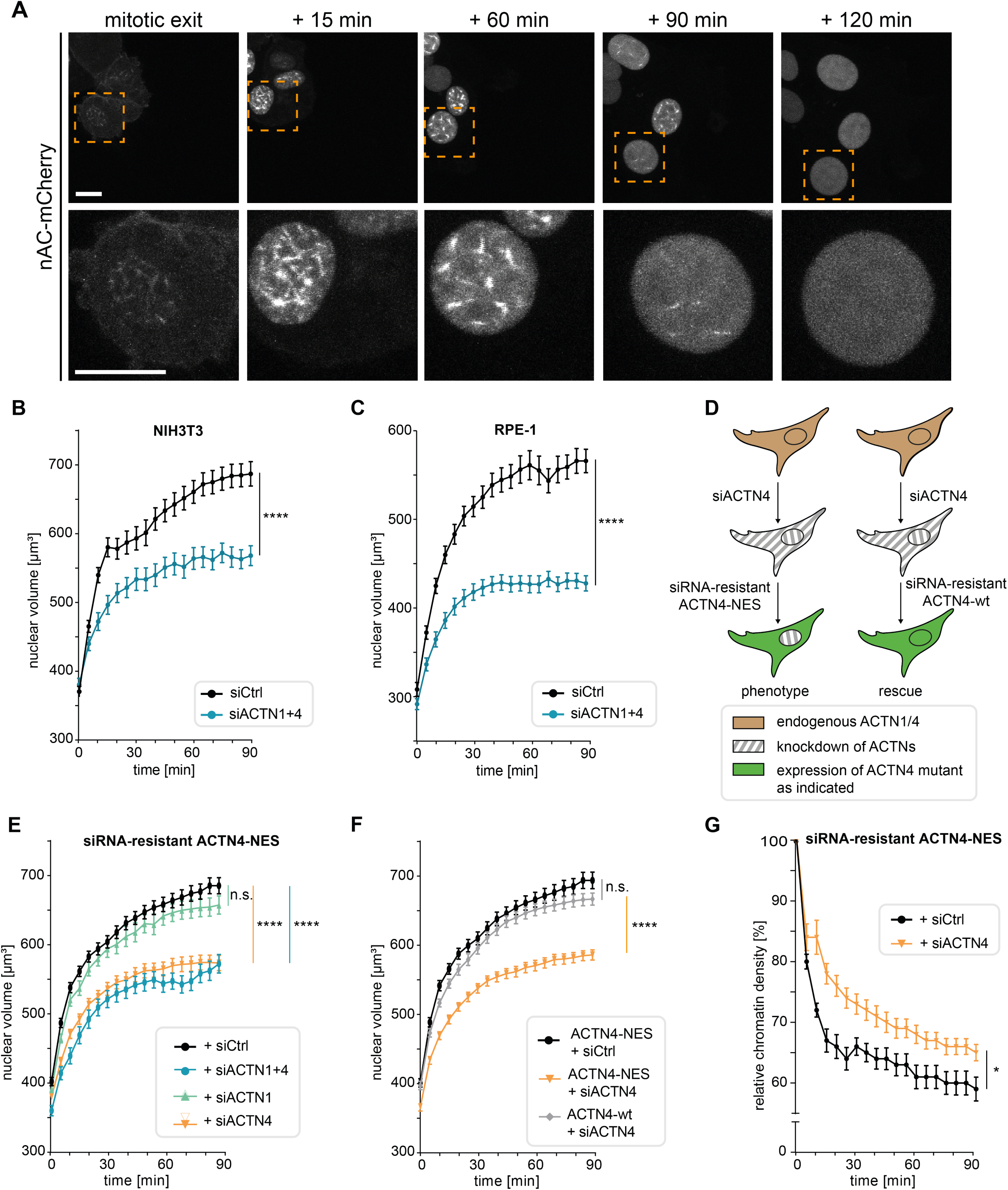
Nuclear ACTN4 is required for nuclear volume expansion after mitotic exit. **(A)** NIH3T3 cells expressing nAC-mCherry were imaged for 120 min. Image frames represent maximum projections of 30 Z planes. Lower panel is a magnification of the indicated area (orange box); scale bars 10 µm. **(B)** NIH3T3 cells stably expressing H2B-mCherry were transfected with siCtrl or siACTN1 + siACTN4. After mitotic exit z-stacks (30 planes) were acquired every 5 min. Nuclei were 3D-reconstructed and volume was measured by Imaris. Data are mean ± SEM from n=3 with 22-27 (siCtrl) and 32-42 (siACTN1+4) analyzed nuclei; p(90 min) < 0.0001. **(C)** RPE-1 cells were analyzed as in B. Data are mean ± SEM from n=5 with 32-37 (siCtrl) and 31-39 (siACTN1+4) analyzed nuclei; p(90 min) < 0.0001. **(D)** Cartoon illustrating experimental design of knockdown and reconstitution experiments in E, F and G to achieve cells lacking ACTN4 specifically in the nuclear compartment (left) versus cells expressing ACTN4-wt. **(E)** NIH3T3 cells stably expressing H2B-mCherry and siRNA-resistant ACTN4-NES were transfected with the indicated siRNAs and analyzed as described in B. Data are mean ± SEM from n=5 with 63-71 (siCtrl), 37-41 (siACTN1+4), 52-58 (siACTN1) and 52-55 (siACTN4) analyzed nuclei; p(siACTN4) < 0.0001; p(siACTN1+4) p < 0.0001 by One Way ANOVA at 90 min. **(F)** NIH3T3 cells stably expressing H2B-mCherry and the indicated ACTN4 mutants were transfected with the indicated siRNAs and analyzed as in B. Data are mean ± SEM from n=3 with 63-69 (ACTN4 wt + siACTN4), 63-71 (ACTN4-NES + siCtrl) and 54-58 (ACTN4-NES + siACTN4) analyzed nuclei; p(ACTN4-NES + siACTN4) < 0.0001 by One Way ANOVA at 90 min. **(G)** NIH3T3 cells expressing siRNA-resistant ACTN4-NES were transfected with siCtrl or siACTN4; chromatin density was calculated from H2B-mCherry fluorescence intensities by total nuclear volumes (intensity/μm^3^) and normalized to t=0 (mitotic exit); Data are mean ± SEM from n=5 with 44-48 (siCtrl) and 44-47 (siACTN4) analyzed nuclei; p (90 min) = 0.0135.

Since silencing of ACTNs could lead to cytoplasmic cytoskeletal defects that might conceal alterations in its nuclear functions, we decided to employ reconstitution studies using siRNA-resistant ACTN4 derivatives fused to a nuclear export signal (NES) versus ACTN4 wild-type (ACTN4-wt) control to specifically study ACTN4 depletion in the nucleus (Fig. 3D). Interestingly, cells silenced for ACTN4 or double-silenced for ACTN1 and ACTN4 that expressed siRNA-resistant ACTN4-NES displayed significantly reduced nuclear volume expansion after mitotic exit (Fig. 3E). Notably, knockdown of ACTN1 alone did not lead to significant reduction in nuclear volume expansion under these conditions (Fig. 3E), indicating that nuclear ACTN4 but not ACTN1 is required for correct volume expansion of daughter cell nuclei after mitotic exit. As a control, we reconstituted ACTN4 knockdown cells by re-expression of siRNA resistant ACTN4-wt, which fully restored nuclear volume expansion (Fig. 3F). This underscores the importance for an ACTN4 function inside the nuclear compartment. Consistent with this notion and in line with our previous findings that nuclear actin assembly promotes chromatin decondensation during nuclear expansion (Baarlink et al., 2017), we observed significantly higher chromatin densities at mitotic exit in ACTN4 silenced cells expressing siRNA-resistant ACTN4-NES (Fig. 3G). Together, these data reveal a specific nuclear function for ACTN4 but not ACTN1 in promoting nuclear expansion after cell division.

To assess the role of ACTN4-mediated actin rearrangement in the nucleus, we disrupted the N-terminal ABD by deleting its calponin-homology 1 (CH1) domain and fused it to a nuclear localization signal (ACTN4ΔCH1-NLS) for nuclear targeting. As expected, ACTN4ΔCH1-NLS showed a predominant nuclear localization (Suppl. Fig. 1A). Notably, NIH3T3 cells expressing ACTN4ΔCH1-NLS displayed significantly reduced nuclear volume expansion after mitotic exit as compared to ACTN4-wt expressing cells (Suppl. Fig. 1B). We therefore decided to use this approach to specifically investigate the impact of nuclear ACTN4 on F-actin formation in early G1. Quantifying F-actin formation is a challenge since a single actin filament measures only 5-7 nm in diameter (Grazi, 1997). In addition, accurate quantification of nAC-mCherry signal is not possible because the actin chromobody labels both G-actin and F-actin, leading to high background signals. We therefore performed live-cell imaging of cells undergoing mitosis followed by subsequent phalloidin-AlexaFluor647 staining to exclusively label F-actin and to perform super-resolution microscopy imaging of the same cells at mitotic exit (Fig. 4A). By direct stochastic optical reconstruction microscopy (dSTORM) imaging (Endesfelder and Heilemann, 2015), we were able to reveal nuclear actin filament formation in cells expressing ACTN4ΔCH1-NLS or ACTN4-wt and to analyze the number and thickness of filament bundles. Interphase cells expressing ACTN4-wt were used to determine the background signal that could either be caused by non-specifically bound phalloidin molecules or by very short F-actin polymers (Fig. 4B). Interestingly, nuclei of ACTN4ΔCH1-NLS cells at early G1 exhibited significantly fewer fluorophore (phalloidin-AF647) localizations than ACTN4-wt cells (Fig. 4C) and this decrease was comparable to interphase cells expressing ACTN4-wt (Fig. 4C). In contrast, nuclei of ACTN4-wt cells fixed in early G1 displayed significantly more fluorophore localizations per nuclear area as well as a significantly higher actin filament number than ACTN4ΔCH1-NLS cells (Fig. 4D), indicating that the ACTN4 derivative lacking the ABD interferes with proper nuclear F-actin formation at mitotic exit. Notably, while we observed fewer filament structures in ACTN4ΔCH1-NLS nuclei, those filaments displayed significantly decreased widths as compared to ACTN4-wt (Fig. 4E). We hence determined the resolution of our dSTORM images to estimate the expected widths of single actin filaments. By measuring the experimental localization precision by NeNA (Endesfelder et al., 2014), which ranged from 10-15 nm in all images, we could approximate our resolution to about 35 nm. Accounting for the actual width of an actin filament of 5-7 nm, the size of phalloidin-AF647 molecules of about 2-3 nm used for labeling the structures (Turkowyd et al., 2016), and the precision of image drift correction of 3-4 nm (Balinovic et al., 2019), we thus expected single filaments to appear at widths of around 45-50 nm in our images. Therefore, we conclude that nuclei of ACTN4ΔCH1-NLS expressing cells mainly exhibited few single actin filaments, whereas the ACTN4-wt expressing cells displayed thicker and more abundant F-actin structures resembling bundled actin filaments (Fig. 4B, D and E).

**Figure 4.**
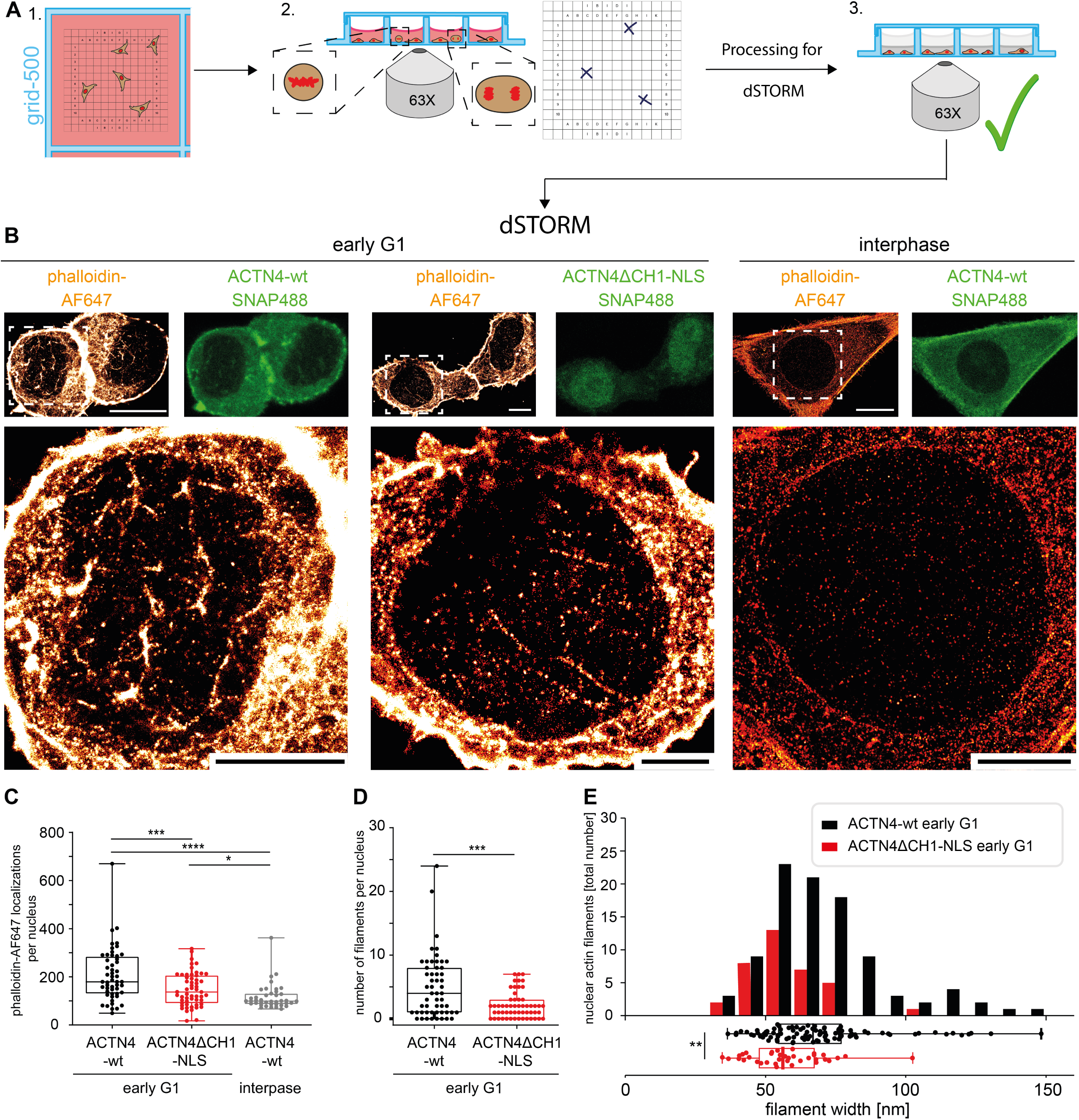
ACTN4 is necessary for postmitotic nuclear actin assembly and actin filament bundling. **(A)** Live imaging in NIH3T3 cells expressing ACTN4 constructs stained with SNAP488 provided positions of mitotic cells. After fixation, cells were stained with phalloidin-AlexaFluor647, rechecked for proper processing and transferred to dSTORM imaging. **(B)** Upper panel shows expression of ACTN4 constructs (green, confocal image) and phalloidin-AF647 staining in the same cells (orange, dSTORM image); lower panel represents zoom of the indicated areas (white boxes); scale bars x µm. **(C)** Phalloidin-AF647 localizations per early G1 nuclei; data shown as mean ± 2.5-97.5 percentile: 52 ACTN4-wt nuclei, 55 ACTN4ΔCH1-NLS nuclei and 40 interphase nuclei (ACTN4-wt) from 5 independent experiments. **(D)** Number of filaments in the same nuclei; data statistics as in (C). **(E)** Actin filament widths for ACTN4-wt and ACTN4ΔCH1-NLS early G1 cells; 96 filaments (ACTN4-wt) and 36 filaments (ACTN4ΔCH1-NLS) from 5 independent experiments.

Here we found that postmitotic nuclear volume expansion is dependent on ACTN4. The underlying mechanism appears to be based on the ability of ACTN4 to bundle actin filaments (Honda et al., 1998; Winkelman et al., 2016), however, we also observed a role for actin assembly by nuclear ACTN4. Interestingly, ACTN4 has been previously shown to affect actin assembly or filament turnover (Kemp and Brieher, 2018). Hence, it is conceivable that a coordinated mechanism between actin assembly and bundling might be at work, further emphasizing the highly complex and dynamic nature of the actin cytoskeleton. It is tempting to speculate that nuclear actin bundles may enable the incorporation of myosin and thus the formation of contractile bundles that could generate forces to allow expansion of the nuclear lamina and chromatin rearrangement. Such hypothesis is further supported by a recent study visualizing single myosin IV molecules moving along intranuclear actin filaments to support long-range chromatin rearrangements (Große-Berkenbusch et al., 2020). Future studies will aim to address the potential role of myosins in postmitotic nuclear expansion and chromatin reorganization.

## Materials and Methods

### Cell culture, transfection and viral transduction

HEK293T, NIH3T3 and RPE-1 were maintained in DMEM (HPSTA – high glucose, stable glutamine and sodium pyruvate; Capricorn Scientific) supplemented with 10% fetal calf serum (FCS; Thermo Fisher Scientific) under standard conditions at 37°C in a 5% CO2 environment. HEK293T cells were transfected using the calcium phosphate method. NIH3T3 and RPE-1 cells were transfected using Lipofectamine 2000 (Thermo Fisher Scientific) following manufacturer’s protocol. Gene silencing was obtained by RNAi. Cells were transfected with siRNAs (FlexiTube, Qiagen) using Lipofectamine RNAiMAX (Thermo Fisher Scientific) following manufacturer’s protocol. Following siRNAs were used: AllStars Negative Control siRNA: AATTCTCCGAACGTGTCACGT; Mm_ACTN1_2: CCGAGTTGATTGACTATGGAA; Mm_ACTN4_5: CAGGGATGGGCTCAAACTTAT; Hs_ACTN1_9: GACCATTATGATTCTCAGCAA and Hs_ACTN4_5: ACGCAGCATCGTGGACTACAA. For virus production, HEK293T cells were transfected using calcium phosphate method with 4-6 µg plasmid DNA and 8 µg of each pMD2.G and psPAX2 (Addgene). After 48 h, growth medium containing lentivirus was harvested and added to freshly seeded target cells. Cells were kept under BSL2 conditions until virus titer was under detection limit. If necessary, cells were subjected to FACS-based cell sorting.

### Molecular cloning

DNA fragments were obtained and amplified by PCR using Phusion Hot Start II High-Fidelity DNA Polymerase (Thermo Fisher Scientific) according to manufacturer’s protocol. Ligation reaction was performed with T4 ligase (Thermo Fisher Scientific) and transformed into DH5α competent bacterial cells. Plasmids were extracted by using the NucleoSpin^®^ Plasmid Miniprep Kit (Macherey-Nagel). All extracted plasmids were sent for DNA sequencing, conducted by Microsynth SeqLab. Deletion mutants and point mutations were created using the conversion extension method (Ito et al., 1991). To allow for nuclear export, the NES sequence of HIV-Rev (LPPLERLTL) was inserted at the C-terminus of ACTN4. Constructs were ligated into pEGFP-N1 (Clontech) or pSNAP-Flag-N1 vector (Baarlink et al., 2017). To create inducible lentiviral constructs, mutants were first subcloned into pEntr11 (Invitrogen) before subcloning into pInducer20 Puro using the LR Clonase™ (Invitrogen), following manufacturer’s instructions. The SV40 large T antigen nuclear localization signal (NLS) PPKKKKRKV was used as previously reported (Plessner et al., 2015) and fused to the ACTN4ΔCH1 mutant. siRNA resistance was obtained by mutating 6 bases (the 3^rd^ of each triplet; silent mutations) in the siRNA recognition site of the RNA. Hs siRNAs 1_7 (5’-AAGGATGATCCACTCACAAAT-3’) and 4_7 (5’-CAGGACATGTTCATCGTCCAT-3’) were used as templates. Constructs have been verified by sequencing (SeqLab) and functionality was proved by expression and simultaneous RNAi (Western Blot and staining). Constructs are also resistant to Mm siRNA 1_2 and 4_5 and Hs siRNA sequences 1_9 and 4_5.

The following primers were used:

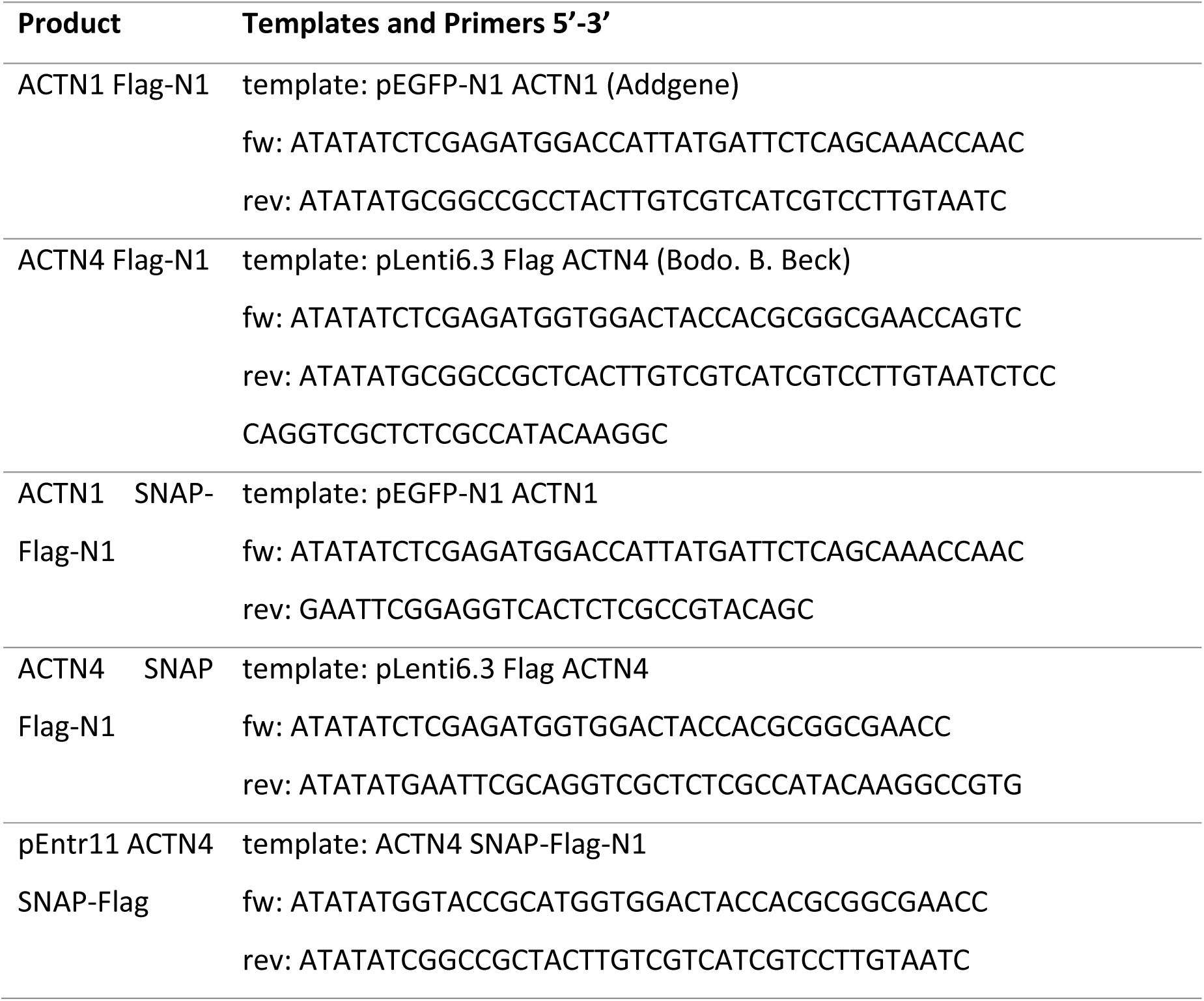
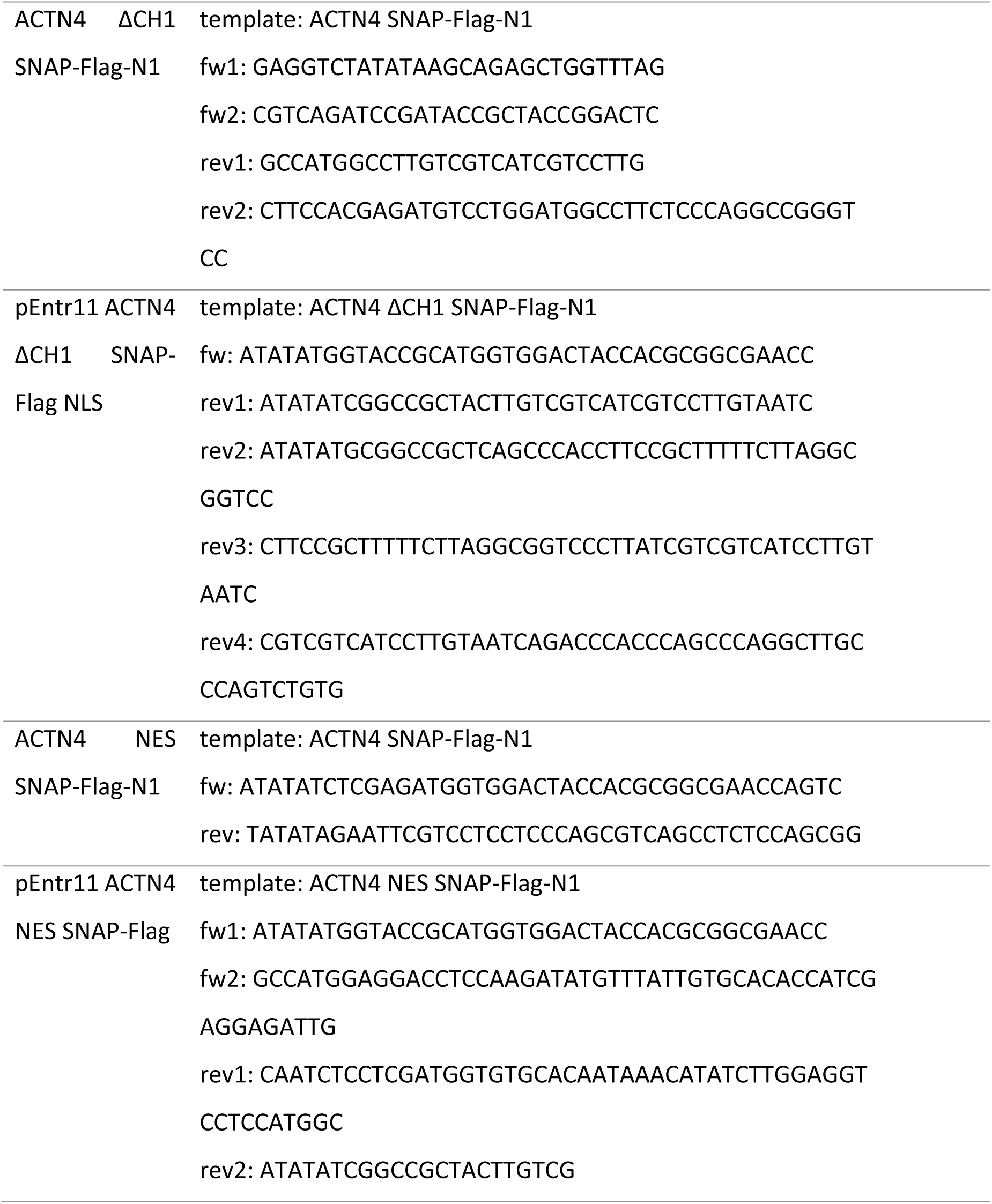

### Real-Time RT-PCR

RNA extraction was performed with TRIzol reagent (Peqlab) according to the manufacturer’s protocol. The reverse transcription was obtained using RevertAid Reverse Transcriptase (Fermentas) and RT-PCR following manufacturer’s instructions. Attained cDNA was quantified in qPCR using iQ™ SYBR^®^ Green super mix (Bio-Rad). The following primers were used: GAPDH fw: 5’-CCCTTCATTGACCTCAACTA-3’; GAPDH rev: 5’-CCAAAGTTGTCATGGATGAC-3’; ACTN1 mm fw: 5’-GACCATTATGATTCCCAGCAGAC-3’; ACTN1 mm rev: 5’-CGGAAGTCCTCTTCGATGTTCTC-3’; ACTN4 mm fw: 5’-ATGGTGGACTACCACGCAG-3’; ACTN4 mm rev: 5’-CAGCCTTCCGAAGATGAGAGT-3’; ACTN1 hs fw: 5’-CAGCGACATCGGTCATCTACATCGG-3’; ACTN1 hs rev: 5’-GTTACACATGGAGGCAGCTCAGGTG-3’; ACTN4 hs fw: 5’-CTGCTGCACTGTGGCTGCTGGAATC-3’; ACTN4 hs rev: 5’-GGCAACCGAGTGGTTCCAGTGGGC-3’. Relative mRNA levels were calculated using the comparative ΔΔCT model (Rao et al., 2013) normalized to GAPDH cDNA, serving as house-keeping gene.

### Microscopy, nuclear volume and chromatin density assays

Images were generated at LSM700 or LSM800, respectively, confocal laser-scanning microscopes (Zeiss), each equipped with a 63X, 1.4 NA oil objective and the ZEN black (for LSM700) or Zen blue (for LSM800) software (Zeiss) essentially as described previously (Grikscheit et al., 2015; Hinojosa et al., 2017). Cells were seeded on glass cover slips, fixed and stained following standard procedures. Antibody labelling was performed with primary anti-ACTN4 (rabbit; ENZO) and secondary anti-rabbit-AlexaFluor488 antibody (goat; Thermo Fisher Scientific). Nuclear compartment was labelled with DAPI. Live cell imaging was performed with 8 well µ-slide ibidi chambers (ibidi) in DMEM with 10 % FCS at 37°C in a CO2-humidified incubation chamber (Pecon, CO2 module S1). Cells expressing H2B-mCherry were used for nuclear volume and chromatin density assays. Images were processed with ZEN blue software and ImageJ/Fiji. Brightness and contrast were optimized, and maximum intensity projections were generated from Z stack images, where indicated. IMARIS software (Bitplane) was used to generate 3D surfaces (from 30 Z planes) based on nuclear-specific signal and to measure respective volumes over time. Chromatin densities were calculated by dividing the sum of H2B mCherry fluorescence intensities by total nuclear volumes (intensity/µm^3^) (Baarlink et al., 2017). Each value was normalized to the value of corresponding timepoint 0. For visualization of ACTN4 dynamics a stable cell line expressing nAC-mCherry and doxycycline-inducible ACTN4-GFP was used. 24h before imaging, ACTN4-GFP expression was induced by 1µg/ml doxycycline. Images were acquired with a CSU-X1 spinning disc confocal microscope (Yokogawa), with 405/488/561/640 laser lines and a photometrics Prime sCMOS camera. Cells were analyzed with a 100x objective at 37°C in a CO2-humidified incubation chamber. ACTN4 dynamics were analyzed by IMARIS tracking using the autoregressive motion algorithm.

### dSTORM sample preparation, imaging and image analysis

Cells were cultured in a grid-500 8 well ibidi (ibidi), imaged with the 63X objective of a Zeiss LSM 700 and positions of cells in metaphase or anaphase were marked. In early G1, cells were fixed in 1 % glutaraldehyde and 0,05 % Triton X-100 in cytoskeleton buffer (CSK) for 1 min at room temperature (RT), followed by a second fixation in 3 % glutaraldehyde in CSK for 10 min at RT and a quenching step in 10 mg/ml NaBH_4_ in water for 10 min. Cells were then blocked with 100 µM L-lysine (Sigma) in ImageIT™ FX Signal Enhancer (Invitrogen) for 1 h at RT and incubated with a 1:50 dilution of phalloidin-AF647 (Thermo Fisher Scientific) for 96 h at 4 °C. After postfixation with 4% formaldehyde in PBS and washing with 0.05 % Tween20 in PBS, IR beads (FluoSphere infrared fluorescent Carboxylate-Modified Microspheres; Thermo Fisher Scientific) were diluted 1:50 in phenol red-free matrigel (Corning; VWR) and added to each well. Matrigel was solidified at 37 °C for 1h and postfixed with 4 % formaldehyde in PBS for 10 min at RT. PBS was added to avoid drying and marked positions were verified by confocal imaging prior to dSTORM.

dSTORM experiments were conducted on a customized automated inverted Ti-Eclipse microscope (Nikon). The imaging procedure was adapted from (Virant et al., 2018). Briefly, the custom build microscope setup contained a CFI Apochromat TIRF 100x oil objective (NA 1.49, Nikon), appropriate dichroic and filters (ZET405/488/561/640 nm dichroic mirror, BrightLine HC 689/23 bandpass, both AHF Analysentechnik), 488 nm and 640 nm lasers (488 nm Sapphire, 640 nm OBIS, both Coherent Inc.) and an emCCD camera iXON ULTRA 888 (Andor) with a pixel size of 129 nm for fluorescence detection. Both lasers were modulated by an acousto optical tunable filter (AOTF, Gooch and Housego). Z-Focus was controlled by a commercial perfect focus system (Nikon). The setup was operated using a customized version of µManager (Edelstein et al., 2014). Before each imaging experiment, the power of the 640 nm laser was adjusted to a final intensity of 1-2 kWcm^-2^ in the sample to ensure consistent conditions for different experimental days. Before imaging, PBS was removed from the wells and dSTORM buffer was added: 100 mM mercaptoethylamine (MEA) with a glucose oxygen scavenger system (van de Linde et al., 2011). The sample was illuminated in HILO (Highly Inclined and Laminated Optical sheet, (Tokunaga et al., 2008) mode and the marked positions were recorded at 20 Hz for 38,000-40,000 frames.

Fluorescent single molecule spots from the image acquisition were localized using ThunderSTORM (Ovesný et al., 2014). The localization files were further processed using customized scripts written in Python programming language (Python Software Foundation, https://www.python.org) and were kindly provided by Dr. Bartosz Turkowyd, Endesfelder group, MPI Marburg, to correct for sample drift during image acquisition using the signals from the 100 nm diameter infrared beads in each sample and to filter out-of-focus signal (80 nm < PSF sigma < 200 nm, uncertainty < 35 nm). Each corrected file was checked for quality by measuring the full width at half-maximum (FWHM) of infrared beads. Drift corrections with a resulting FWHM of 70 nm and larger were revisited. From the processed localization files, the final experimental localization precision was determined by calculating the NeNA precision value (Endesfelder et al., 2014). Super resolution images were reconstructed with a pixel size of 10 nm using RapidSTORM 3.0 (Wolter et al., 2012) and were processed with a Gaussian blur according to their individual NeNA localization precision. The number of localizations per nucleus was measured by selecting the nuclear region with the software swift (written in C++, Endesfelder group, unpublished) and normalized to the nuclear area. Filament widths were measured with a customized script for Fiji (Schindelin et al., 2012) where filaments were manually selected by a segmented line profile (Virant et al., 2018). To minimize errors due to selection and pixelation, selected areas were shifted 0.5 pixels (corresponding to 5 nm) in all directions. From this, five measurements (ROIs) were attained which were further straightened to remove filament curvatures. After projection of the filament ROIs along their long axis, each profile was fitted by a Gaussian distribution. Resulting values show the full filament width at half-maximum value. Histogram bin size was determined based on the Freedman-Diaconis rule (Freedman and Diaconis, 1981). Number of filaments was determined with the script that has also been used for filament width measurements. The number of analyzed filaments was related to each daughter nucleus.

### Subcellular fractionation, Co-Immunoprecipitation and F-actin pulldown

For cell cycle experiments, cells were arrested by adding 10 µM of the CDK1 inhibitor RO3306 (Merck) for 18 h before washed out and release (2h) to conduct experiments when early G1 was reached. Subcellular fractionations and phalloidin-based pull-downs were performed as described previously (Baarlink et al., 2017). Briefly, cells were pelleted and resuspended before vortexing and centrifugation. The supernatant was used to obtain the cytoplasmic extract. The pelleted nuclei were extracted in lysis buffer before sonication and centrifugation to obtain the nucleoplasmic fraction. Immunoblotting was performed with α-tubulin as cytoplasmic and histone H3 (Cell Signaling) as nucleoplasmic marker to verify successful fractionation. For co-immunoprecipitations, agarose beads conjugated with Flag-M2 antibody (Sigma-Aldrich) were equilibrated in lysis buffer before subcellular fractionations were loaded onto the beads at 4°C for 90 min. Beads were resuspended processed for Western analysis. Phalloidin pulldown assays were performed with streptavidin magnetic Dynabeads (Thermo Fisher Scientific) and Biotin-Phalloidin (Thermo Fisher Scientific).

### Statistics

Statistical analysis was performed using GraphPad Prism 7. Data is presented as mean ± SEM. Statistical significance was evaluated with Two-Way ANOVA for multiple comparisons or unpaired two-tailed Student’s *t* tests for comparison of two groups (as indicated in the figures). Statistical significance are indicated as p ≤ 0.05: *, p ≤ 0.05; **, p ≤ 0.01; ***, p ≤ 0.001; and ****, p < 0.0001.

## Acknowledgements

We thank the laboratory members for helpful discussions, Bodo Beck for providing the pLenti6.3 Flag-ACTN4 plasmid and Bartosz Turkowyd for providing some Python scripts for dSTORM analysis. We further thank Hartmann Raifer (BMFZ, University of Marburg) for FACS analysis and Dominique Brandt for technical assistance. Work in the laboratory of R.G. is funded by the DFG, under Germany’s Excellence Strategy (EXC-2189, project ID: 390939984) and the HFSP program (grant ID: RGP0021/2016). Work in the laboratory of U.E. was supported by the Max Planck Society and by SYNMIKRO.

## Figure Legends

**Suppl. Fig. 1.**
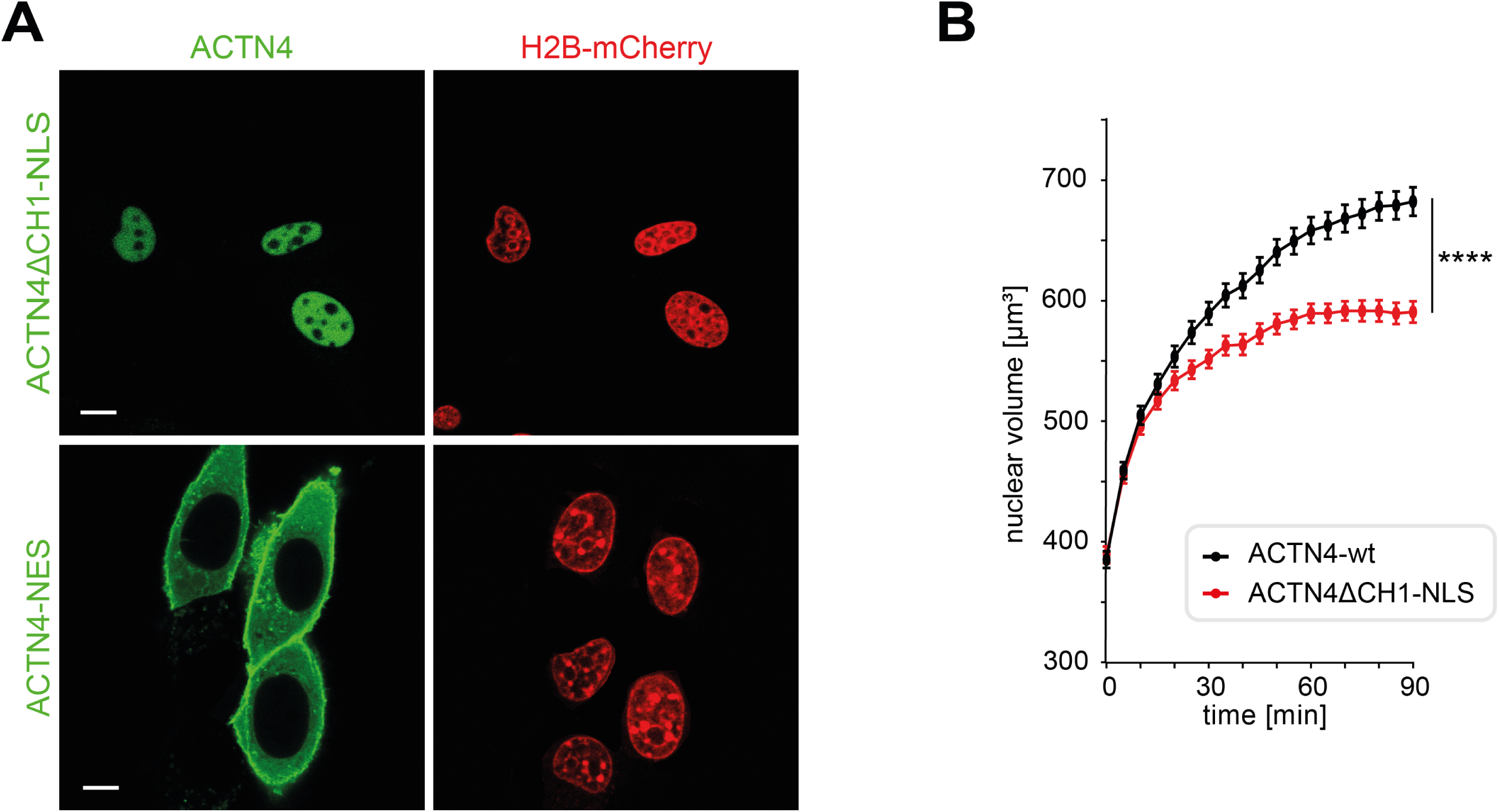
**(A)** NIH3T3 cells expressing H2B-mCherry and the indicated ACTN4 constructs were stained by SNAP488; scale bars 10 µm. **(B)** NIH3T3 cells stably expressing H2B-mCherry were transfected with ACTN4-wt and ACTN4ΔCH1-NLS. After mitotic exit z-stacks (30 planes) were acquired, every 5 min. Nuclei were 3D-reconstructed and volume was measured by Imaris software. Data are shown as mean ± SEM from n=4 with 56-62 nuclei (ACTN4-wt) and 51-57 nuclei (ACTN4ΔCH1-NLS); p < 0.0001.

**Movie 1**

NIH3T3 cells stably expressing nAC-mCherry and doxycycline-inducible ACTN4-snap were analyzed by time-lapse microscopy in early G1. ACTN4-snap was labeled by SiR 647 (green). Arrows label dynamic ACTN4 clusters. Scale bar 5µm. The movie sequence is also part of Fig 2A.

**Movie 2**

Dividing NIH3T3 cells stably expressing nAC-mCherry were analyzed by time-lapse microscopy. The movie is a 3D-reconstruction by imaris software. Scale bar 5µm. The movie sequence is also part of Fig. 3A.

